# Angiotensin-(3–4) modulates angiotensin converting enzyme 2 (ACE2) downregulation in proximal tubule cells due to overweight and undernutrition: implications regarding the severity of renal lesions in Covid-19 infection

**DOI:** 10.1101/2020.06.29.178293

**Authors:** Rafael Luzes, Humberto Muzi-Filho, Amaury Pereira-Acácio, Thuany Crisóstomo, Adalberto Vieyra

## Abstract

The renal lesions – including severe acute kidney injury – are severe outcomes in SARS-CoV-2 infections. There are no reports regarding the influence of the nutritional status on the severity and progress of these lesions. Ageing is also an important risk factor. In the present communication we compare the influence of overweight and undernutrition in the levels of renal angiotensin converting enzymes 1 and 2. Since the renin-angiotensin-aldosterone system (RAAS) has been implicated in the progress of kidney failure during Covid-19, we also investigated the influence of Angiotensin-(3–4) (Ang-(3–4)) the shortest angiotensin-derived peptide, which is considered the physiological antagonist of several angiotensin II effects. We found that both overweight and undernutrition downregulate the levels of angiotensin converting enzyme 2 (ACE2) without influence on the levels of ACE1 in kidney rats. Administration of Ang-(3–4) recovers the control levels of ACE2 in overweight but not in undernourished rats. We conclude that chronic and opposite nutritional conditions play a central role in the pathophysiology of renal Covid-19 lesions, and that the role of RAAS is also different in overweight and undernutrition.

## Introduction

Since the first report of Covid-19 in China on 31 December 2019 and isolation of the SARS-CoV-2 virus on January 7, 2020, the number of infections is still growing with an accelerated rate, as well as the number of deaths worldwide (1). Considering all countries and regions, the number of infections reached >9 million people and more than 475,000 deaths on June 24, 2020 [2]. Even though the illness was considered a viral infection targeting the respiratory system [3], some cases developed into septic shock [4,5], in which acute kidney injury (AKI) plays a central role [6]. Age, cancer, cardiovascular diseases and metabolic diseases – such as diabetes – are currently considered the main risk factors for the explosive outbreak of the pandemic [7].

With respect to the kidney and AKI, it is important to remember that angiotensin converting enzyme 2 (ACE2), considered to be the way of entrance of SARS-CoV-2 into cells [8,9], is highly expressed in the membranes of proximal tubule cells [10], the tubular segment specially affected in AKI [11]. Moreover, levels of ACE2 seem to be important during Covid-19 infections, and this has been recently reviewed in terms of an apparent paradox demonstrating friend-and-foe roles in the evolution of the viral infection [12]. The levels of ACE2 decrease with ageing in different tissues [13,14], but there are no reports regarding the influence of the nutritional status on these levels, especially in obesity and undernutrition, which, as recently emphasized, must be taken into account to investigate the impact of Covid-19 [15].

## Methods

The protocols were approved by the Committee for Ethics in Animal Experimentation from the Federal University of Rio de Janeiro (#101/16 and #012/19). In this letter we report the effect of a chronic administration of 2 different diets on the levels of ACE2 in the external portion of rat renal cortex (*cortex corticis*), where <95% of the cell population comprises the proximal tubules [16]. One was a deficient diet that provokes undernutrition, mimicking the alimentary habits of different regions from developing countries, the so-called Regional Basic Diet (RBD) [17], which is characterized by low protein content (also of poor quality), associated with low lipid content, and absence of vitamins and mineral supplementation. The other diet had high lipid and low carbohydrate content, with an average of 50% higher NaCl than the control (CTR) diet [18] (HL diet), that leads to obesity with an elevated deposition of visceral fat, a tissue with high levels of inflammatory factors [19] that can stimulate the renin-angiotensin-aldosterone system (RAAS) [20,21]. The experiments were conducted during the development of 2 projects in our laboratory during the last year, and we believe that the results – if presented together to allow comparisons – could shed some light on the mechanisms underlying the progression and the severity of renal lesions due to Covid-19.

Male Wistar rats were fed: (i) with the RBD from weaning (28 days of age) until 90 days of age; (ii): with the HL diet from 56 to 162 days of age. The CTR groups received a commercial chow for rodents. At the end of the periods of exposure to the different diets, the rats received Ang-(3–4), one oral dose (80 mg/kg body mass) 24 h before sacrifice in the undernutrition study, and 4 doses at 12 h intervals from 48 h before sacrifice. Angiotensin-(3–4) (Ang-(3–4), Val-Tyr), the shortest angiotensins-derived peptide [22,23], can be viewed as a physiological antagonist of Angiotensin II (Ang II) effects [24]. An enriched plasma membrane fraction from proximal tubule cells was obtained as previously described [25]. Western blotting analyses were carried out as described in the figure legends.

## Results and Discussion

Figure 1 shows that ACE2 levels decrease with age in CTR rats, in contrast to the outcome with ACE. Juvenile rats are the CTR animals from the undernutrition study and, according to Quinn [26], 90 days of age correspond to 21 human years and 162 days (adult) correspond to 38 human years. The 50% decrease in ACE2 in this short period of life (Figure 1A,B) represents, *per se*, an increase in the risk of kidney proximal tubules damage – and therefore of AKI – during SARS-CoV-2 infection in normonourished rats, and this could also be applicable to human kidneys, as in the lungs [12]. Figure 1A,C shows that ACE levels remained unmodified over that relatively long period of life and, therefore, this could contribute to the ACE/ACE2 imbalance that would worsen kidney injuries, as is the case with lung and heart [27], especially in the presence of comorbidities or modifications in the renal local RAAS.

**Figure 1.**
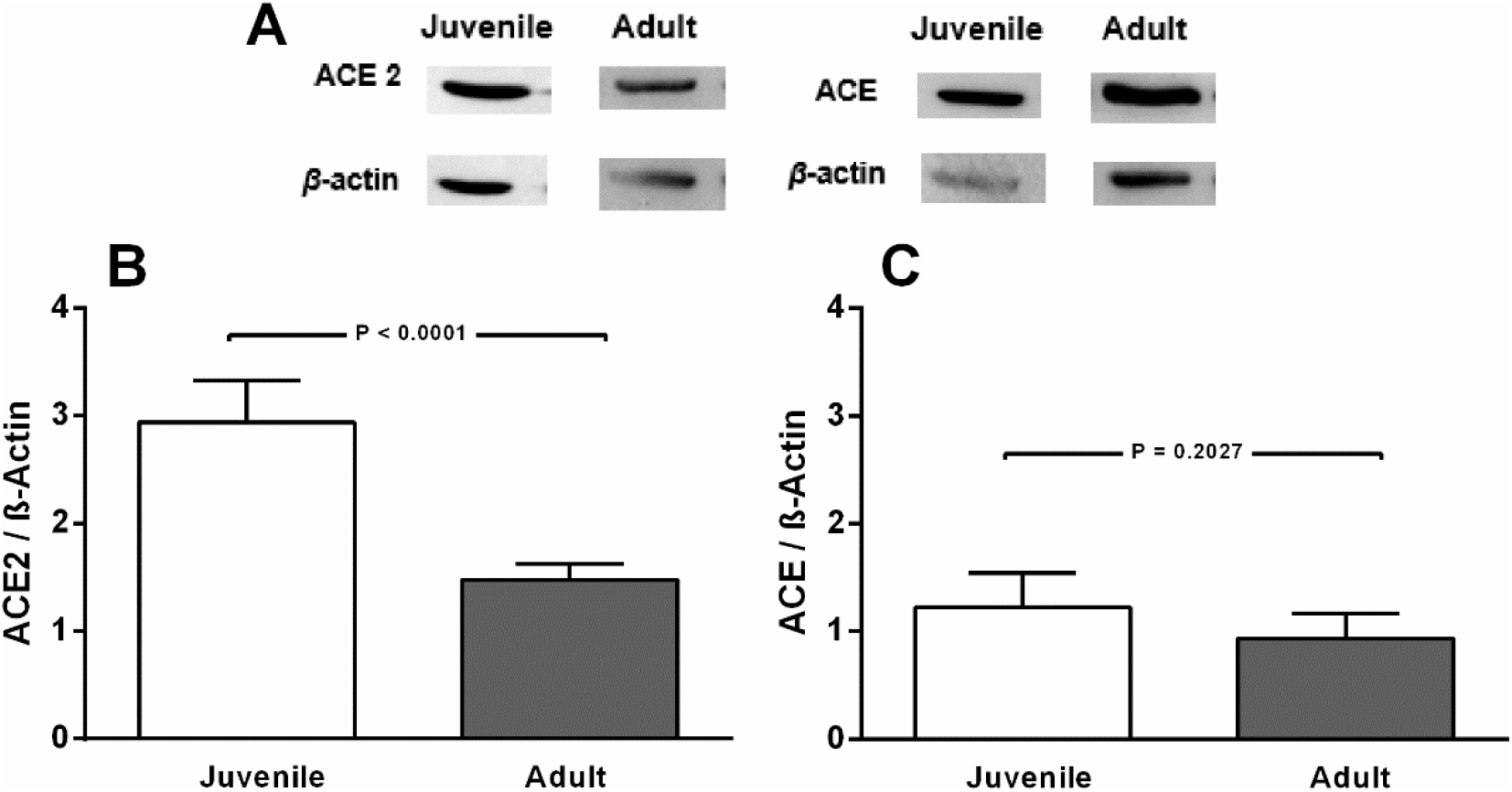
ACE2, but not ACE levels decrease in normonourished CTR rats aged 162 days (adult) compared to CTR 90 days-old rats (juvenile). (A) Representative immunoblottings from renal *cortex corticis* of juvenile and adult rats, with β-actin as the loading control. Bargraph representation of ACE2 (B) and ACE (C) levels in renal *cortex corticis* of juvenile and adult rats. Bars indicate means ± SEM (n = 4−8 different membrane preparations). Statistical differences were estimated by unpaired Student’s *t*-test. P values are indicated within the panels. Differences were set at P<0.05. In A, the representative blots from juvenile and adult rats were the same for ACE2 and ACE. Cutting was unavoidable because different studies are being compared. Loading controls were run on the same blot.

In the next figures our results show that modifications in the nutritional status profoundly modified ACE2 abundance in the renal *cortex corticis*, which is also modulated in a different manner by antagonizing RAS in cases of overweight or undernutrition. Figure 2A,B shows that the abundance of ACE2 decreased 30% in the HL (overweight) rats whose body mass was ~20% higher than that of the CTR group. Administration of Ang-(3–4) to the overweight rats increased by 80 and 150%, when the ACE2 levels are compared with those in the CTR and HL rats, respectively, with no influence in the normonourished group. This observation supports the view that, in overweight animals, antagonizing the Ang II/type 1 Ang II receptor (AT_1_R) axis by Ang-(3–4) could avoid adverse renal injury in SARS-CoV-2 infections. In addition, this last finding implies that, beyond upregulation of the Ang II/Ang-(1–7)/Mas axis [28], intrarenal RAAS can be counteracted – in the case of obesity/overweight – by the end product of a pathway that involves progressively shorter Ang II-derived peptides, as we demonstrated a decade ago [29]. The mechanism underlying the effect of the peptide could be an action as “allosteric enhancer” that induces a second binding site in AT_2_R with a very high affinity for Ang II [30], possibly involved in the modulation of ACE2 formation. Another possibility would be a beneficial formation of heterodimers involving these modified AT_2_R and Mas receptors, and that are able to act on the ACE2, as in the case of blood pressure [31]. That Ang-(3–4)-mediated upregulation of renal ACE2 occurred in overweight rats, but not in the CTR group, is indicative that a “pro-hypertensive tissular microenvironment” (high Ang II) develops in animals fed with a diet rich in lipids, causing downregulation of ACE2 (Figure 2A,B) and kidney injury during a SARS-CoV-2 attack. The levels of ACE (Figure 2A,C) remained unmodified either by diet or Ang-(3–4) treatment.

**Figure 2.**
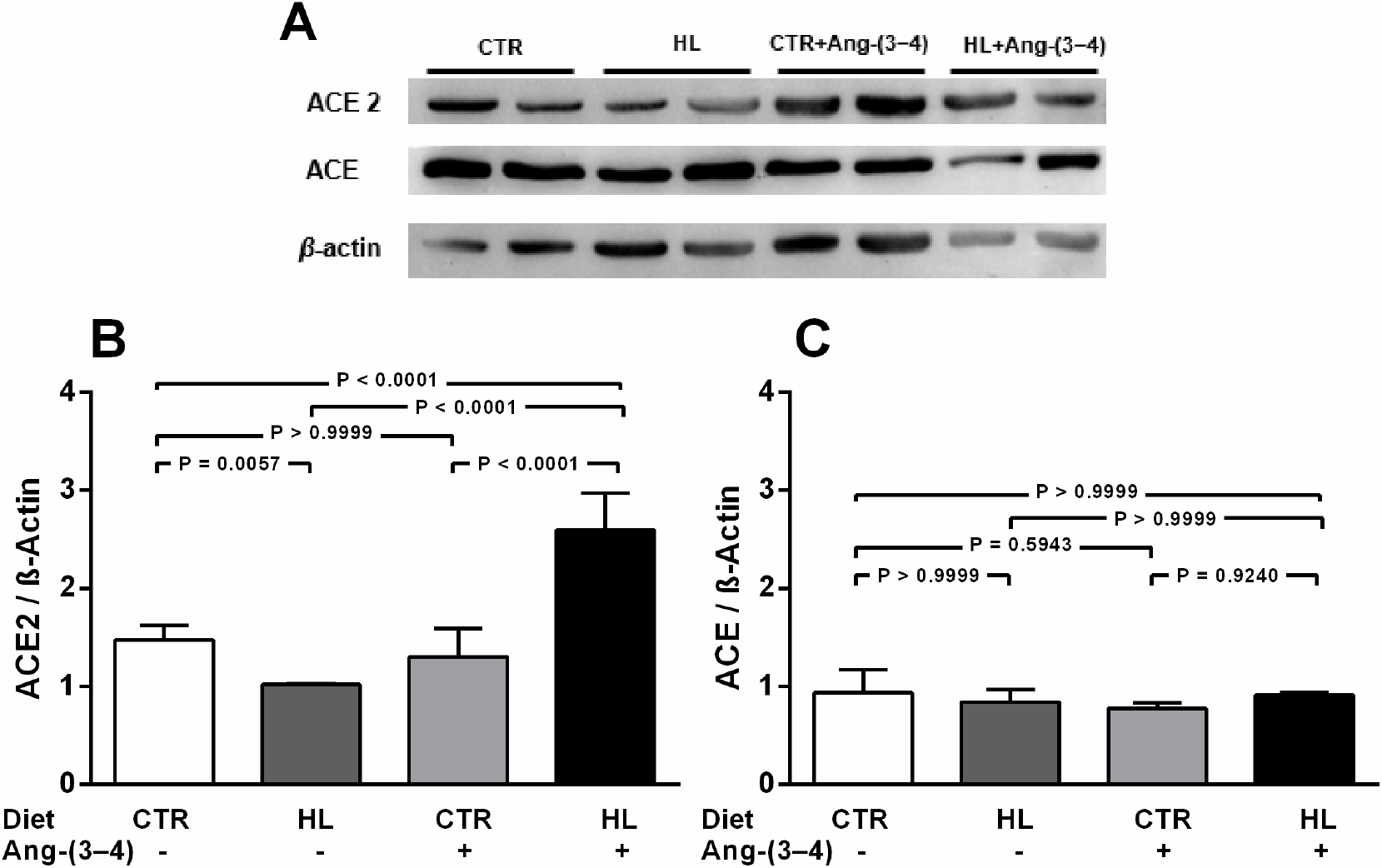
Overweight downregulates ACE2, but not ACE, in adult rats on a high lipid (HL) diet. Effects of Ang-(3–4). (A) Representative immunoblottings from renal *cortex corticis* of control and overweight adult rats; β-actin was used as the loading control. Bargraph representation of ACE2 (B) and ACE (C) levels in renal *cortex corticis*. Bars indicate means ± SEM (n = 4−8 different membrane preparations). Combinations of diets and Ang-(3–4) administration as indicated on the *abscissae*. Statistical differences were estimated using one-way ANOVA followed by Bonferroni’s test for the selected pairs indicated within the panels. Significant differences were set at P<0.05. In A, the figure shows bands from the same gel.

The influence of chronic undernutrition on renal ACE2 levels present some similarity, but also a huge difference, compared with overweight rats (Figure 3). The similarity is in the downregulation of the enzyme level by RBD alone (~40%) (Figure 3A,B), i.e. a value that did not differ from the overweight-induced downregulation seen in Figure 2A,B (~35%). Again, it may be that activation of the Ang II/AT_1_R axis now in a “pro-hypertensive tissular microenvironment” induced by the multideficient diet contributes for the notable decrease in ACE2, and, since there was no modification in ACE (Figure 3A,C), to the important ACE/ACE2 imbalance in proximal tubules that might be crucial in the kidney, as seems to be the case in the lung [27]. On this account, we have strong evidence for the existence of a “pro-hypertensive tissular microenvironment” in chronically undernourished rats: the increased number of Ang II-positive cells in the tubulointerstitium [32] leads to an increase of the local RAAS. In contrast with that observed in the case of overweight rats, the administration of Ang-(3– (i) did not recover (or upregulates) ACE2 in undernourished rats, and (ii) provoked a similar decrease in CTR rats. The second observation meets with the supposition that AT_2_R acts in parallel with AT_1_R in promoting tissular damage in kidney tissue (recently reviewed in [27]). There is a possibility of an adjuvant and independent AT_2_R-associated pathway that, being altered by an amino acid imbalance [33] due to undernutrition, downregulates ACE2 synthesis and levels. A second possibility is the formation of heterodimers between AT_1_R and abnormal AT_2_R, which could be modulated by Ang II and Ang-(3–4), as in the case of spontaneously hypertensive rats [34]. As noted above, these rats are of a juvenile age. Therefore, undernutrition could favor severe kidney lesions in young people affected by Covid-19, an issue that has not been explored until the present.

**Figure 3.**
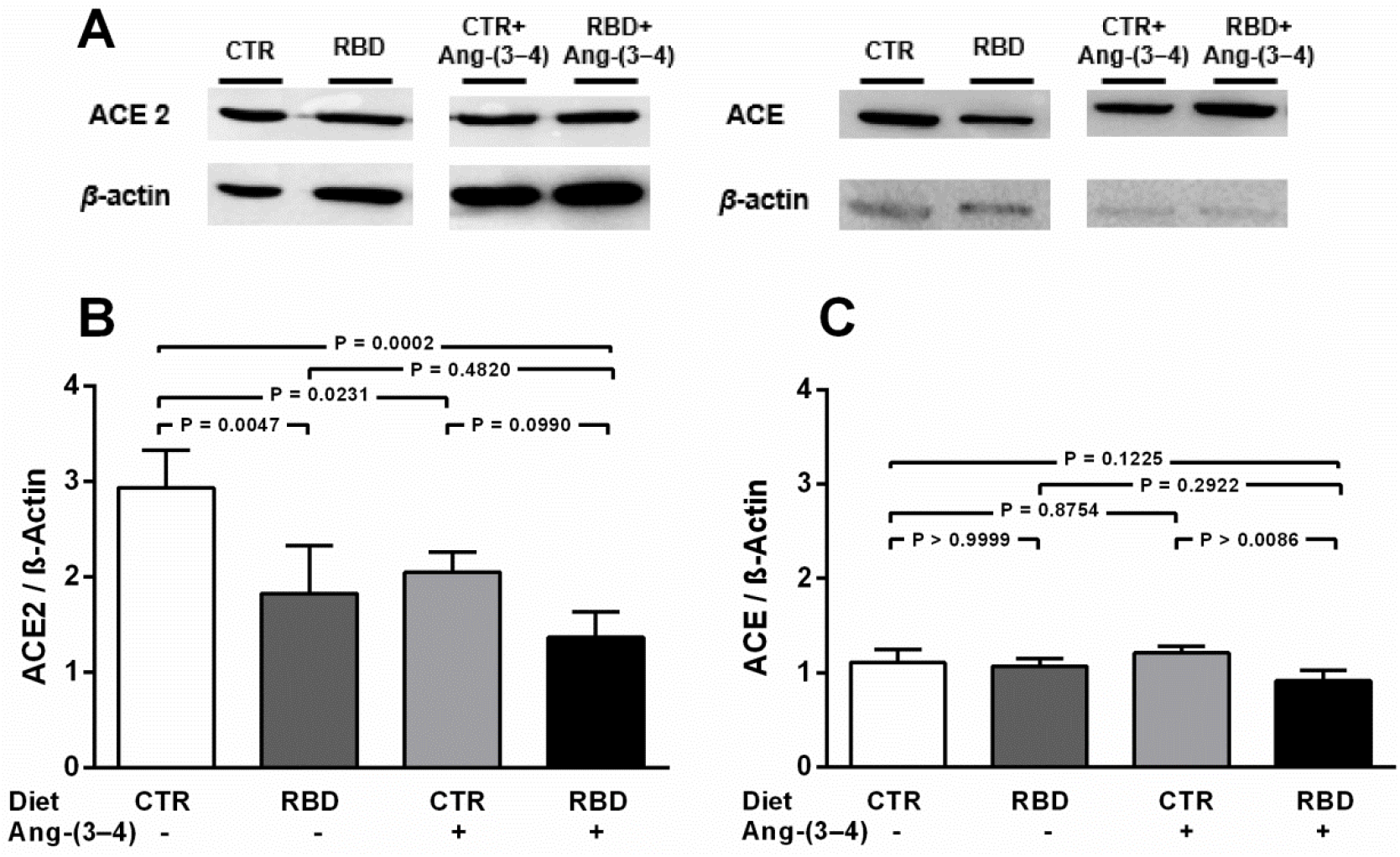
Chronic undernutrition downregulates ACE2, but not ACE, levels in juvenile rats. Effects of Ang-(3–4). (A) Representative immunoblottings from renal *cortex corticis* of juvenile rats; β-actin was used as loading control. Bargraph representation of ACE2 (B) and ACE (C) levels in renal *cortex corticis*. Bars indicate means ± SEM (n = 4 different membrane preparations). Combinations of diets and Ang-(3–4) administration as indicated on the *abscissae*. Statistical differences were estimated using one-way ANOVA followed by Bonferroni’s test for the selected pairs indicated within the panels. Significant differences were set at P<0.05. In A, the representative immunoblottings were derived from the same gel, and were cut to remove information irrelevant to the work described in this letter. Loading controls were run on the same blot.

## Conclusion

In summary, our evidence is that overweight and undernutrition favor downregulation of the ACE2 level in proximal tubule cells, being a threat during the clinical evolution of the SARS-CoV-2 infection. Moreover, the results demonstrate that counteracting RAAS by Ang-(3–4) administration influences ACE2 synthesis in an opposite way in overweight and undernourished rats.

## Acknowledgments

The excellent technical assistance of Glória Costa-Sarmento and Danilo Bezerra is acknowledged. The authors also acknowledge the English corrections by BioMedES UK.

## References

[1] World Health Organization. Novel Coronavirus – China. January 2020. Available at: https://www.who.int/csr/don/12-january-2020-novel-coronavirus-china/en/. Accessed June 24, 2020.

[2] European Centre for Disease Prevention and Control. COVID-19 situation update worldwide. Available at: https://www.ecdc.europa.eu/en/geographical-distribution-2019-ncov-cases. Accessed June 24, 2020.

[3] Xie P, Ma W, Tang H, Liu D. Severe COVID-19: A review of recent progress with a look toward the future. Front Public Health 8 (2020) 189. https://doi.org/10.3389/fpubh.2020.00189.

[4] Hantoushzadeh S, Norooznezhad AH. Possible cause of inflammatory storm and septic shock in patients diagnosed with (COVID-19). Arch Med Res 51 (2020) 347–348. https://doi.org/10.1016/j.arcmed.2020.03.015.

[5] Robba C, Battaglini D, Pelosi P, Rocco PRM. Multiple organ dysfunction in SARS-CoV-2: MODS-CoV-2. Expert Rev Respir Med (2020) Online ahead of print. https://doi.org/10.1080/17476348.2020.1778470.

[6] Perico L, Benigni A, Remuzzi G. Should COVID-19 concern nephrologists? Why and to what extent? The emerging impasse of angiotensin blockade. Nephron 144 (2020) 213–221. https://doi.org/10.1159/000507305.

[7] Jordan RE, Adab P, Cheng KK. Covid-19: Risk factors for severe disease and death. BMJ 368 (2020) m1198. https://doi.org/10.1136/bmj.m1198.

[8] Hoffmann M, Kleine-Weber H, Schroeder S, Krüger N, Herrler T, Erichsen S, Schiergens TS, Herrler G, Wu NH, Nitsche A, Müller MA, Drosten C, Pöhlmann S. SARS-CoV-2 cell entry depends on ACE2 and TMPRSS2 and is blocked by a clinically proven protease inhibitor. Cell 181 (2020) 271–280.e8. https://doi.org/10.1016/j.cell.2020.02.052.

[9] Walls AC, Park YJ, Alejandra Tortorici M, Wall A, McGuire AT, Veesler D. Structure, function, and antigenicity of the SARS-CoV-2 spike glycoprotein. Cell 181 (2020) 281–292.e6. https://doi.org/10.1016/j.cell.2020.02.058.

[10] Ye M, Wysocki J, William J, Soler MJ, Cokic I, Batlle D. Glomerular localization and expression of angiotensin-converting enzyme 2 and angiotensin-converting enzyme: Implications for albuminuria in diabetes. J Am Soc Nephrol 17 (2006) 3067–3075. https://doi.org/10.1681/ASN.2006050423.

[11] Bianco M, Lopes JA, Beiral HJV, Filho JDD, Frankenfeld SP, Fortunato RS, Gattass CR, Vieyra A, Takiya CM. The contralateral kidney presents with impaired mitochondrial functions and disrupted redox homeostasis after 14 days of unilateral ureteral obstruction in mice. PLoS One 14 (2019) e0218986. https://doi.org/10.1371/journal.pone.0218986.

[12] Verdecchia P, Cavallini C, Spanevello A, Angeli F. The pivotal link between ACE2 deficiency and SARS-CoV-2 infection. Eur J Intern Med 76 (2020) 14–20. https://doi.org/10.1016/j.ejim.2020.04.037.

[13] Xie X, Chen J, Wang X, Zhang F, Liu Y. Age- and gender-related difference of ACE2 expression in rat lung. Life Sci 78 (2006) 2166–2171. https://doi.org/10.1016/j.lfs.2005.09.038.

[14] Yoon HE, Kim EN, Kim MY, Lim JH, Jang IA, Ban TH, Shin SJ, Park CW, Chang YS, Choi BS. Age-associated changes in the vascular renin-angiotensin system in mice. Oxid Med Cell Longev 2016 (2016) 6731093. https://doi.org/10.1155/2016/6731093.

[15] Puig-Domingo M, Marazuela M, Giustina A. COVID-19 and endocrine diseases. A statement from the European Society of Endocrinology. Endocrine 68 (2020) 2–5. https://doi.org/10.1007/s12020-020-02294-5.

[16] Whittembury G, Proverbio F. Two modes of Na extrusion in cells from guinea pig kidney cortex slices. Pflügers Arch 316 (1970) 1–25.

[17] Teodósio NR, Lago ES, Romani SA, Guedes RC. A regional basic diet from northeast Brazil as a dietary model of experimental malnutrition. Arch Latinoam Nutr 40 (1990) 533–547.

[18] Reeves PG. Components of the AIN-93 diets as improvements in the AIN-76A diet. J Nutr 127 (1997) 838S–841S. https://doi.org/10.1093/jn/127.5.838S.

[19] Costa RM, Neves KB, Tostes RC, Lobato NS. Perivascular adipose tissue as a relevant fat depot for cardiovascular risk in obesity. Front Physiol 9 (2018) 253. https://doi.org/10.3389/fphys.2018.00253.

[20] Francischetti EA, Genelhu VA. Obesity-hypertension: An ongoing pandemic. Int J Clin Pract 61 (2007) 269–280. https://doi.org/10.1111/j.1742-1241.2006.01262.x.

[21] Hall JE, do Carmo JM, da Silva AA, Wang Z, Hall ME. Obesity-induced hypertension: interaction of neurohumoral and renal mechanisms. Circ Res 116 (2015) 991–1006. https://doi.org/10.1161/CIRCRESAHA.116.305697.

[22] Matsufuji H, Matsui T, Seki E, Osajima K, Nakashima M, Osajima Y. Angiotensin I-converting enzyme inhibitory peptides in an alkaline protease hydrolyzate derived from sardine muscle. Biosci Biotechnol Biochem 58 (1994) 2244–2245. https://doi.org/10.1271/bbb.58.2244.

[23] Saito Y, Wanezaki K, Kawato A, Imayasu S. Antihypertensive effects of peptide in sake and its by-products on spontaneously hypertensive rats. Biosci Biotechnol Biochem 58 (1994) 812–816. https://doi.org/10.1271/bbb.58.812.

[24] Dias J, Axelband F, Lara LS, Muzi-Filho H, Vieyra A. Is angiotensin-(3–4) (Val-Tyr), the shortest angiotensin II-derived peptide, opening new vistas on the renin-angiotensin system? J Renin Angiotensin Aldosterone Syst 18 (2017) 1470320316689338. https://doi.org/10.1177/1470320316689338.

[25] Vieyra A, Nachbin L, de Dios-Abad E, Goldfeld M, Meyer-Fernandes JR, de Moraes L. Comparison between calcium transport and adenosine triphosphatase activity in membrane vesicles derived from rabbit kidney proximal tubules. J Biol Chem 261 (1986) 4247–4255.

[26] Quinn R. Comparing rat’s to human’s age: How old is my rat in people years? Nutrition 21 (2005) 775–777. https://doi.org/10.1016/j.nut.2005.04.002.

[27] Sriram K, Insel PA. A hypothesis for pathobiology and treatment of COVID-19: The centrality of ACE1/ACE2 imbalance. Br J Pharmacol (2020) Online ahead of print. https://doi.org/10.1111/bph.15082.

[28] Santos RAS, Simões e Silva AC, Maric C, Silva DMR, Machado RP, Buhr I, Heringer-Walther S, Pinheiro SVB, Lopes MT, Bader M, Mendes EP, Lemos VS, Campagnole-Santos MJ, Schultheiss HP, Speth R, Walther T. Angiotensin-(1–7) is an endogenous ligand for the G protein-coupled receptor MAS. Proc Natl Acad Soc USA 100 (2003) 8258–8263. https://doi.org/10.1073/pnas.1432869100.

[29] Axelband F, Dias J, Miranda F, Ferrão FM, Barros NM, Carmona AK, Lara LS, Vieyra A. A scrutiny of the biochemical pathways from Ang II to Ang-(3–4) in renal basolateral membranes. Regul Pept 158 (2009) 47–56. https://doi.org/10.1016/j.regpep.2009.08.004.

[30] Axelband F, Dias J, Miranda F, Ferrão FM, Reis RI, Costa-Neto CM, Lara LS, Vieyra A. Angiotensin-(3–4) counteracts the Angiotensin II inhibitory action on renal Ca^2+^-ATPase through a cAMP/PKA pathway. Regul Pept 177 (2012) 27–34. https://doi.org/10.1016/j.regpep.2012.04.004.

[31] Patel S, Hussain T. Dimerization of AT_2_ and Mas Receptors in Control of Blood Pressure. Curr Hypertens Rep 20 (2018) 41. https://doi.org/10.1007/s11906-018-0845-3.

[32] Silva PA, Muzi-Filho H, Pereira-Acácio A, Dias J, Martins JFS, Landim-Vieira M, Verdoorn KS, Lara LS, Vieira-Filho LD, Cabral EV, Paixão ADO, Vieyra A. Altered signaling pathways linked to angiotensin II underpin the upregulation of renal Na^+^-ATPase in chronically undernourished rats. Biochim Biophys Acta 1842 (2014) 2357–2366. https://doi.org/10.1016/j.bbadis.2014.09.017.

[33] Harper AE, Benevenga NJ, Wohlhueter RM. Effects of ingestion of disproportionate amounts of amino acids. Physiol Rev 50 (1970) 428–558. https://doi.org/10.1152/physrev.1970.50.3.428.

[34] Dias J, Ferrão FM, Axelband F, Carmona AK, Lara LS, Vieyra A. ANG-(3‒4) inhibits renal Na^+^-ATPase in hypertensive rats through a mechanism that involves dissociation of ANG II receptors, heterodimers, and PKA. Am J Physiol Renal Physiol 306 (2014) F855–F863. https://doi.org/10.1152/ajprenal.00488.2013.

